# Unravelling microalgal-bacterial interactions in aquatic ecosystems through 16S rRNA gene-based co-occurrence networks

**DOI:** 10.1101/2022.01.05.475147

**Authors:** B.L.D. Uthpala Pushpakumara, Kshitij Tandon, Anusuya Willis, Heroen Verbruggen

## Abstract

Interactions between microalgae and bacteria can directly influence the global biogeochemical cycles but the majority of such interactions remain unknown. 16S rRNA gene-based co-occurrence networks have potential to help identify microalgal-bacterial interactions. Here, we used data from 10 Earth microbiome projects to identify potential microalgal-bacterial associations in aquatic ecosystems. A high degree of clustering was observed in microalgal-bacterial modules, indicating densely connected neighbourhoods. *Proteobacteria* and *Bacteroidetes* predominantly co-occurred with microalgae and represented hubs of most modules. Our results also indicated that species-specificity may be a global characteristic of microalgal associated microbiomes. Several previously known associations were recovered from our network modules, validating that biologically meaningful results can be inferred using this approach. A range of previously unknown associations were recognised such as co-occurrences of *Bacillariophyta* with uncultured *Planctomycetes OM190* and *Deltaproteobacteria* order *NB1-j. Planctomycetes* and *Verrucomicrobia* were identified as key associates of microalgae due to their frequent co-occurrences with several microalgal taxa. Despite no clear taxonomic pattern, bacterial associates appeared functionally similar across different environments. To summarise, we demonstrated the potential of 16S rRNA gene-based co-occurrence networks as a hypothesis-generating framework to guide more focused research on microalgal-bacterial associations.

## Introduction

Microalgae (photosynthetic micro-eukaryotes) and bacteria are the most widespread and dominant planktonic organisms in aquatic ecosystems. As primary producers, microalgae form the foundations of the food web and directly influence global carbon and nutrient cycling, as well as the energy flow in aquatic ecosystems. A wide spectrum of associations between microalgae and bacteria have been reported, predominantly related to nutrient exchange, with important bottom-up effects on primary production. Associations include increased bioavailability of vitamins (e.g.B12^1–3^), metals (e.g. iron^4,5^) and growth promoting hormones^6^ by bacteria in exchange for organic carbon from the algae. Apart from many beneficial interactions, some bacteria can have negative influences on algae by being algicidal and opportunistic pathogens^7–9^. A recent study on the bloom-forming *Phaeocystis* algae revealed that its microbiomes can take both symbiotic and opportunistic modes^10^. Thus, algal-bacterial interactions can range from mutualistic to parasitic, and a complex array of interactions can be hypothesised. For instance, previous work suggests that strong associations may exist between communities of microalgae and bacteria over a large geographical range in the open ocean^11^. However, the knowledge we have so far about specific microalgal-bacterial interactions represents only a small fraction of what potentially occurs in nature. Microalgae are highly diverse and innumerable, therefore gathering knowledge on their associations with equally ubiquitous and diverse bacteria is challenging.

High-throughput DNA sequencing such as 16S and 18S rRNA gene metabarcoding has been used to deliver insights into bacterial and eukaryotic community composition of diverse ecosystems. Microbial community abundance data can be used to identify associations between community members via microbial association networks^12^. Furthermore, combining community data with environmental data can reveal novel connections between microbial communities and their environment^13^. A network consists of nodes representing taxa and edges representing the associations between taxa. Microbial association networks targeting positive correlations are termed co-occurrence networks. A positive relationship is presumed when taxonomically relevant units co-occur or exhibit similarity in their compositions over multiple samples^14^ and biologically meaningful groups or communities of a network can be identified with network clustering^15^. Also, network metrics such as node degree, closeness centrality for example, can be used for quantitative description of communities and identify ecologically important taxa^16,17^. Various network analyses have reported co-occurring microbial taxa in different environments, identified clusters of microbes representing metabolic consortia, defined keystone species, documented recurrent microbial modules and helped elucidate microbial dark matter in microbial communities^18–21^. Ecological processes governing community structure, such as niche filtering and habitat preference, are thought to be reflected in co-occurrence network modules^22^. Overall, networks provide insights into the structure of a community and taxon co-occurrences, offering perspectives to use them to generate hypotheses about interactions that can be investigated with more focused studies. Thus, environmental sequencing data combined with network analysis can be a convenient addition to the toolkit for deriving potential interactions between organisms across myriads of environments.

Studies targeting 16S rRNA marker genes are generally focused on the bacterial communities and often disregard the chloroplast sequences that are also amplified. The shared origin of the chloroplasts of all oxygenic photosynthetic micro-eukaryotes and cyanobacteria^23^ enable the use of 16S rRNA marker gene to concurrently estimate relative abundances of microalgae and bacteria across samples, facilitating the construction of correlation networks to identify co-occurring taxa and speculate on their potential interactions. Robust co-occurrences can be detected through analysis of a sufficient number of samples, ideally covering temporal and spatial gradients, as these provide adequate variability in taxon abundances^12^. This can be coupled with stringent network building steps such as filtration of low prevalence organisms to reveal significant co-occurrences^24^. The ever-expanding sequence databases can be a great starting point to investigate such correlations. Analysis of the Tara Ocean dataset based on interaction networks has shown that abiotic factors are incomplete predictors of community structure^22^. Similar observations have been shown in phytoplankton blooms where environmental parameters were insignificant in influencing the community structure of plankton^25^. These results indicate that biological interactions are more influential in determining the community structure than environmental factors. Thus, taxon-taxon co-occurrence networks built on taxon compositions alone could capture potential interactions in nature. Our work is based on the premise that previously unknown associations between microalgae and bacteria may be discovered at large scales from existing 16S rRNA gene-based metabarcoding datasets using co-occurrence networks. In this study, we generated taxon-taxon co-occurrence networks using ten 16S rRNA gene metabarcoding datasets from the Earth Microbiome Project (EMP) that have sampled aquatic environments (both marine and freshwaters). These datasets were individually analysed to create local networks and network modules were used to identify significant co-occurrences of microalgae and bacteria.

## Methods

### Data Acquisition

Publicly available EMP datasets^26^ were screened using the Qiita portal^27^ and 10 studies (4 marine and 6 Freshwater environments) targeting aquatic environments were chosen (Figure S1). Since the goal of this study was to individually analyse samples representing a particular environment to generate local networks and reveal co-occurrences, studies targeting any aquatic samples (water and sediment samples) were selected. To our knowledge, based on study and sample information provided on qiita portal for the selected studies, no size fractionation was carried out on the samples (water samples) provided to EMP. The data analysed in this study have amplified the same 16S rRNA gene V4 hypervariable region using 515F-‘GTGCCAGCMGCCGCGGTAA’ and 806R-‘GGACTACHVGGGTWTCTAAT’ primer pair^28,29^ and sequenced on Illumina Hiseq2000. Detailed information on the selected EMP projects can be found in Supplementary material (Table S1) and can also be further explored using Qiita ids provided using the Qiita portal.

### Sequencing Data Workflow

Raw demultiplexed reads were downloaded using the EBI accessions (provided in Qiita portal) from the European Nucleotide Archive and analysed individually employing a single bioinformatic pipeline built on Qiime2 version 2019.10^30^. Primer sequences attached to all reads were trimmed using cutadapt^31^. Sequence denoising, chimera checking and dereplication was performed in DADA2 ^32^ to correct sequencing errors and remove low quality bases. Reads were truncated based on a median quality score of 30. The final outputs of DADA2, an abundance table of Amplicon Sequence Variants (ASVs) and fasta sequences of the ASVs were further processed as described hereafter. Taxonomy was assigned against Silva v132 [30] 16S rRNA gene sequences trained with a Naive Bayes classifier^33^. Chloroplast sequences were filtered using the Silva taxonomy file by identifying sequences assigned as “chloroplast”. These chloroplast sequences were then classified again using a Qiime2-compatible version of PhytoRef^34^ database accessed at^35^. All the algal taxonomic identities with PhytoRef had a confidence score of 0.7 or higher (Table S2 and S3). It is important to note that, each algal node in the networks (see below for details) represents a single ASV, likely representing one algal species or strain in the environment.

The DADA2 abundance table was filtered to create a bacteria-only abundance table (by removing mitochondrial, Archaeal and chloroplast ASVs) and a chloroplast-only abundance table (by retaining only the ASV ids assigned as chloroplast). The bacteria-only table was then collapsed at the genus level to reduce weak links in downstream network building^36^ and was merged with the chloroplast-only table to create the final abundance table. ASVs identified as microalgae were not collapsed since the 16S rRNA gene marker is highly conserved in algae and ASVs likely correspond to species or higher-level taxa^34^. This final abundance table was filtered to remove low prevalence organisms present in less than 10% of samples to prevent them from introducing artefacts in network inference.

### Correlation analysis and network construction

We used a compositional data analysis tool, FastSpar^37^, which is a rapid and parallelizable implementation of the SparCC algorithm^38^ to compute correlations. FastSpar quantifies the correlation between all ASVs and assesses the statistical significance (p-values) of the inferred correlations using a bootstrap procedure. The correlation and p-value matrices created by FastSpar were used to create a network of significant correlations for each study separately. Co-occurrence networks were created using the igraph package^39^ in R studio version 1.4.1106. Undirected weighted networks were created using statistically (p < 0.05) significant correlations > 0.5 or higher (0.5 for freshwater and 0.6 for marine datasets). Coefficient cut-off of 0.5 was selected for freshwater datasets as a value above that resulted in fewer algal nodes in networks compared to 0.5. The idea behind the coefficient cut-off selection was to preserve as many links as possible with microalgae in resulting networks while choosing a considerably higher coefficient cut off to report stronger links. Networks were visualised in Cytoscape version 3.8^40^.

### Network Analysis and module detection

The Network Analyzer plugin^41^ in Cytoscape was used to compute global network properties of each network to get an overview of node specific and edge-specific attributes. Networks were checked for the scale free nature by identifying the presence of highly connected nodes coexisting with nodes with fewer links^42^ using the node attributes. In order to identify communities, networks were clustered using the Cytoscape plugin clusterMaker^43^ using the MCL clustering algorithm^44^ with an inflation value of 2.0. Co-occurring microalgae and bacteria were identified using resulting modules (modules with nodes > or = 4). These co-occurring taxa (identified beyond the taxonomic rank Phylum) were recorded for each environment and summarised in heatmaps using Pheatmap v1.0.12^45^ in R.

## Results and Discussion

### 16S rRNA gene-based co-occurrence networks can recover microalgal-bacterial associations

We generated 10 co-occurrences networks representing aquatic environments using publicly available 16S rRNA gene datasets from the EMP. We reduced noise and false positives by including ASVs present in at least 10% of samples, filtering statistically insignificant and weaker correlations. We also used network modules that are indicative of ecological communities to identify potential interactions. In each co-occurrence network generated, we observed highly connected nodes coexisting with nodes with fewer links. This indicates the scale-free nature of the networks, a characteristic in real world networks^46,47^. The clustered networks (modules) comprised nodes representing microalgae or bacteria and the edges between them were instances of significant co-occurrences.

Analysis of the microalgal-bacterial modules identified 40 algal nodes co-occurring with at least one of 76 bacterial nodes in marine environments and 112 microalgal nodes with at least one of 311 bacterial nodes in freshwater environments (refer Table S2 and S3 for recovered co-occurrences with the correlation values). These significant co-occurrences inferred in marine and freshwater environments are summarised in Figures 1 and 2, respectively. We identified algal nodes at different taxonomic levels although many could not be classified at lower taxonomic levels such as genus or species. To have consistency in algal node annotations, we have only used class-level classification in figures. Tables S2 and S3 provide full taxonomic affiliations of the nodes where possible. Even though taxonomic assignments were made at higher rank, our ASVs represent taxa at lower taxonomic levels. As the breadth of species included in reference databases such as PhytoRef increases, it is likely that more algal ASVs can be assigned to the species-level.

**Figure 1:**
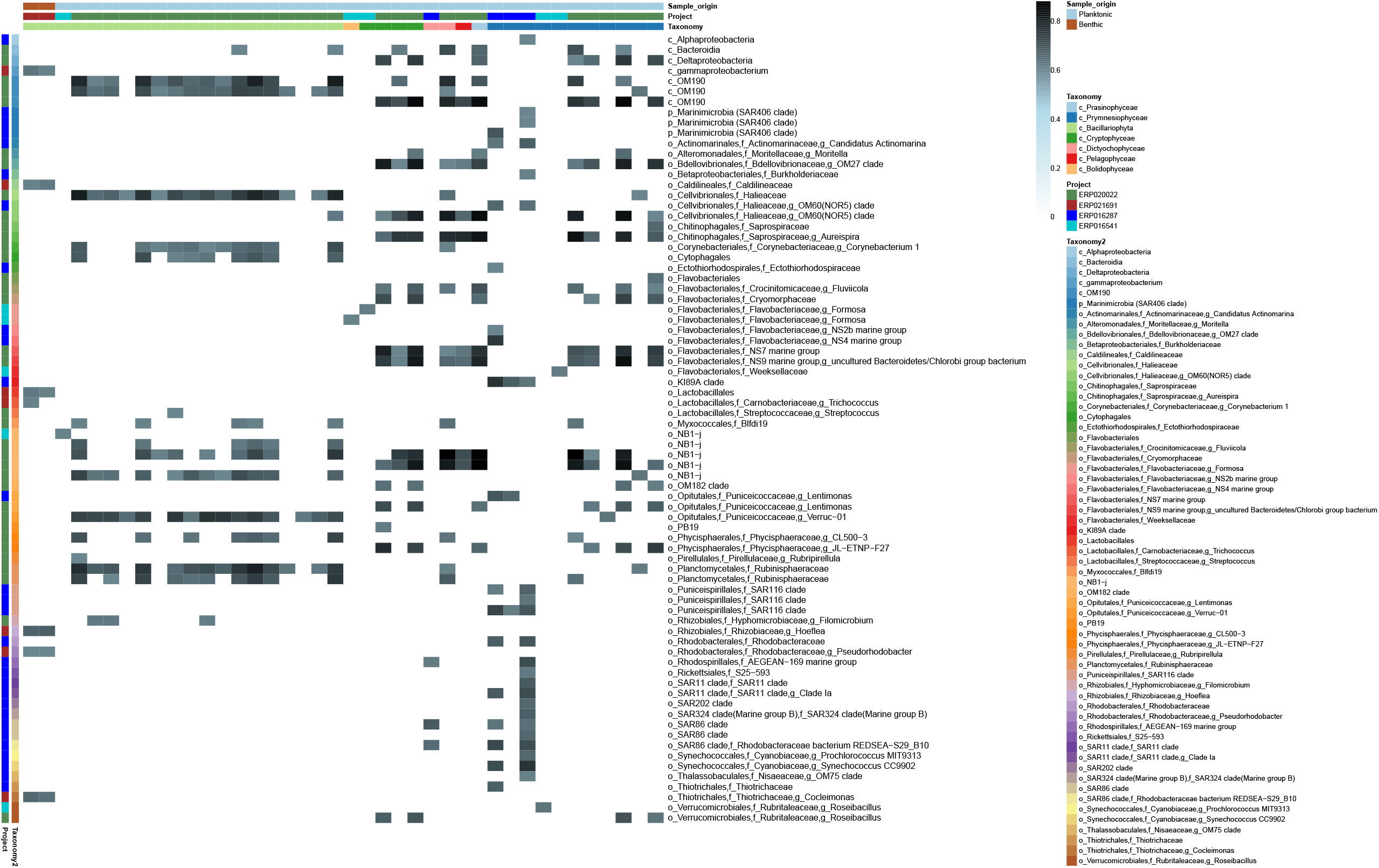
Summarised interactions in marine environments. Correlation heatmap of co-occurring microalgae (columns) and bacteria (rows) in marine environments. “Taxonomy” represents the taxonomic affiliations of microalgal nodes at class level. “Taxonomy2” represents the taxonomic affiliations of bacteria (p_, c_, o_, f_ and g_ represent Phylum, Class, Order, Family and Genus, respectively). “Project” represents the accession numbers of EMP projects and indicate which project the microalgal and bacterial nodes were recovered from. “Sample_origin” indicates if each sample is planktonic or benthic. Heatmap colour gradient indicates correlation coefficients. Refer Table S2 for raw data used to generate the heatmap. Heatmap was generated using Pheatmap v1.0.12 (https://rdrr.io/cran/pheatmap/).

**Figure 2:**
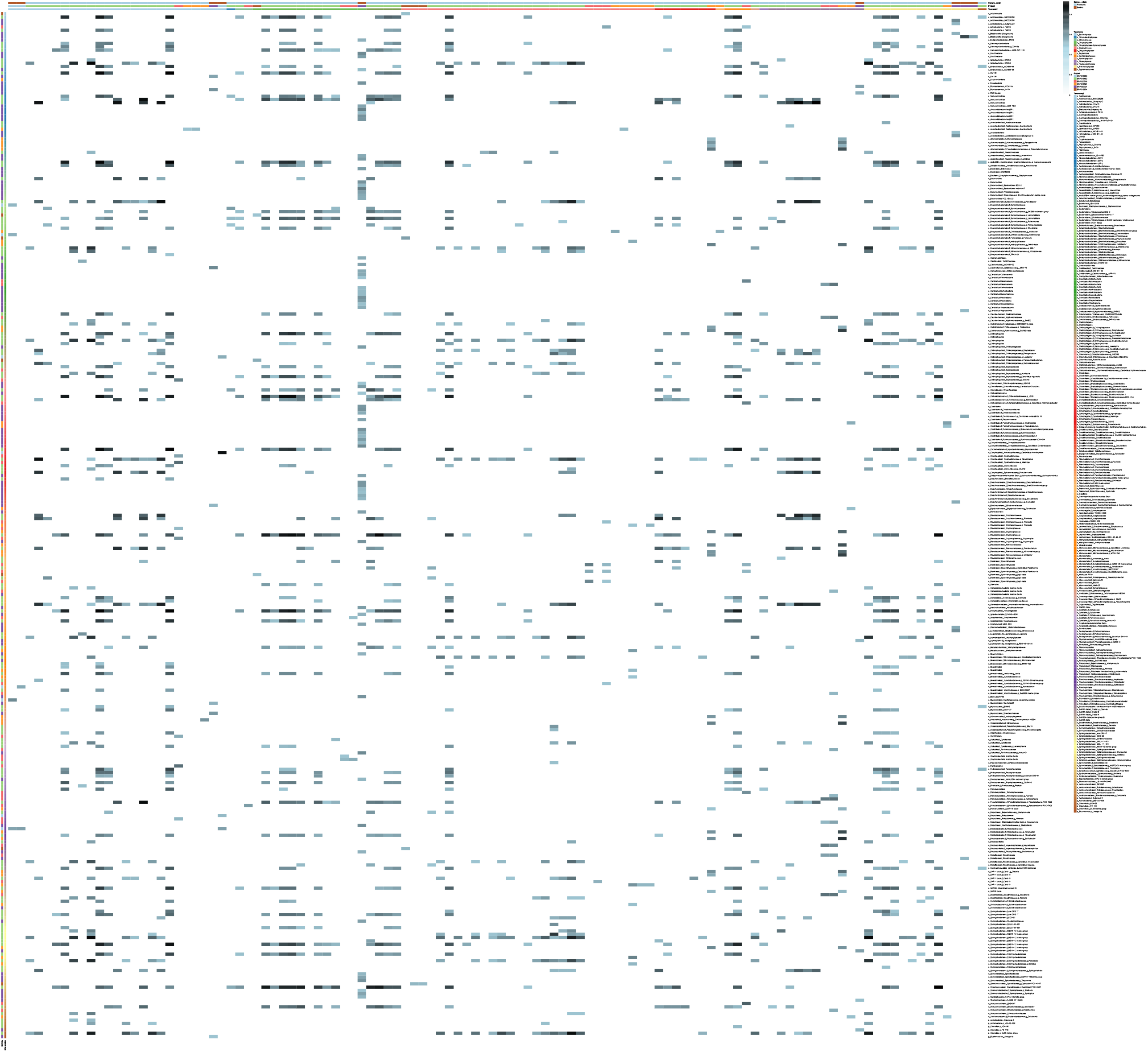
Summarised interactions in Freshwater environments. Correlation heatmap of co-occurring microalgae (columns) and bacteria (rows) in freshwater environments. “Taxonomy” represents the taxonomic affiliations of microalgal nodes at class level. “Taxonomy2” represents the taxonomic affiliations of bacteria (p_, c_, o_, f_ and g_ represent Phylum, Class, Order, Family and Genus, respectively). “Project” represents the accessions of EMP projects and indicate which project the microalgal and bacterial nodes were recovered from. “Sample_origin” indicates if each sample is planktonic or benthic. Heatmap colour gradient indicates correlation coefficients. Refer Table S3 for raw data used to generate the heatmap. Heatmap was generated using Pheatmap v1.0.12 (https://rdrr.io/cran/pheatmap/).

Most modules exhibited high clustering coefficients (> 0.5) indicating that the neighbourhood of microalgal-bacterial communities are generally densely connected. High clustering coefficient values in networks were previously suggested as indicative of cross-feeding relationships and enriched degradation pathways^48^. We identified potential interactions of bacteria with diverse phyla such as *Cryptophyta, Ochrophyta, Haptophyta, Chlorophyta, Streptophyta* and *Euglenophta* representing 17 different taxonomic classes. Bacterial nodes in the marine modules were predominantly represented by *Proteobacteria* and *Bacteroidetes* followed by *Planctomycetes and Verrucomicrobia*. Similar to marine environments *Proteobacteria and Bacteroidetes* dominated the freshwater modules while *Actinobacteria* and *Verrucomicrobia* were the 2nd and 3rd most common bacterial taxa associated with microalgae (Figure S2). We also identified hub nodes with the highest node degree (number of edges connected to a node). Hub nodes represent highly connected nodes and are usually considered as keystone species^49^. Hub nodes (considering the top ten hubs) in both marine and freshwater modules were mostly represented by *Alphaproteobacteria, Gammaproteobacteria* (p_*Proteobacteria*) and *Bacteroidia* (p_*Bacteroidetes*). Other than these, members of *Planctomycetes* (mostly belonging to c_*Planctomycetacia*) were commonly found among the top hub nodes in freshwater modules. High prevalence and their characteristic associations with microalgae may explain the presence of *Proteobacteria* and *Bacteroidetes* as hub nodes in most modules^50,51^. The role of *Planctomycetes* in global nitrogen, carbon and sulphur cycles is gaining attention^52,53^. Thus, microalgal-*Planctomycetes* associations may be playing a crucial role in the global environmental cycles.

Based on the summarised significant co-occurrences, benthic diatoms exhibited different, and fewer, associations compared to planktonic diatoms (Figures 1 and 2). For example, bacterial taxa such as unclassified *Gammaproteobacteria, Caldilineaceae* (*Chloroflexi*), *Trichococcus* (*Firmicutes*), *Hoeflea* (*Proteobacteria*), *Pseudorhodobacter* (*Proteobacteria*) and *Cocleimonas* (*Proteobacteria)* exhibited associations only with marine benthic diatoms and not with marine planktonic diatoms.

We investigated whether this network approach has recovered any known associations identified and confirmed earlier. We recognized known bacterial associates of microalgae such as *Flavobacteriales*^54,55^, *Rhizobiales*^56,57^, *Cytophagales*^58,59^ and *Rhodobacterales*^60^. Previously reported associations of microalgal genera with specific bacterial groups were also recovered in the analysis, including that between *Bacteroidetes* and the cosmopolitan *Prymnesiophyceae* genus *Phaeocystis*^61–63^ (Table S2) and that of diatoms (*Bacillariophyta*) with the genus *Flavobacterium* (Table S3)^64–66^. The ability to identify previously known associations suggests that the 16S rRNA gene-based network approach used in this study can yield biologically meaningful results.

We also compared co-occurrence networks built in our study with previously published networks of these data. Previous analysis of lake Mendota samples (EBI accession: ERP016591) by Kara et al^67^ described characteristics of bacterioplankton co-occurrence networks across three seasons. The network properties of these bacterioplankton networks such as clustering coefficient (0.167 - 0.256) and the characteristic path length (3.27 - 3.88) were comparatively lower than those of our microalgal-bacterial network (clustering coefficient > 0.5, characteristic path length = 3.965). Similar to the observation by Gilbert et al^68^, the co-occurrence network built on Western English Channel samples (EBI accession: ERP016541) featured many *Alphaproteobacteria and Rhodobacterales* nodes.

### Both Intrinsic and Extrinsic factors may influence microalgal-bacterial associations

No clear taxonomic pattern was observed among the co-occurring microalgae and bacteria. In accordance with previous work on phytoplankton-bacteria co-occurrences^67^, taxonomically diverse bacteria co-occurred with different microalgal groups. Moreover, except for a few, there was no recurrence of bacterial genera (the lowest taxonomic identity of bacterial nodes in a module) across modules generated in each project. Since each project represents a specific environment, these observations indicate that each environment harbours its specific co-occurrence relationships. Project specificity in inferred co-occurrences can clearly be seen in the generated heatmaps (Figures 1 and 2). Interestingly, a recent study^68^ has provided evidence for biogeographic differentiation of algal microbiomes, showing ecological boundaries driven by differences in environmental conditions altering the spatial scaling of the algal microbiomes. Another plausible explanation for such observations is the species specificity of algal microbiomes. Although some microalgal nodes could not be taxonomically identified at species level, each microalgal node in a module is an ASV which likely represents a single microalgal species or a strain. Species specificity of algal microbiomes has been predominantly shown using algal cultures^50^. In addition to supporting previous observations, our results show that species-specificity may be a global characteristic of microalgal microbiomes.

To understand if there is any factor driving general patterns of interactions, some insights may be gleaned from known functions of the co-occurring bacteria. Our observation was that despite their taxonomic differences, bacterial associates in modules often shared functional similarities. For instance, microalgae co-occurred with bacterial groups equipped with specific metabolic functional potentials such as algal polysaccharide degradation and provision of vitamins. A few examples of these are, co-occurrences with taxonomically diverse members of *Bacteroidetes*^63^ and *Verrucomicrobia*^69^ with polysaccharide degradation ability and *Rhizobiales, Rhodobacterales* ^70^ and *SAR116* ^71–73^ with Vitamin B12 synthesis. Therefore, our results stemming from observations across multiple datasets suggest that, irrespective of the environment, microalgae are associated with key functional types of bacteria. As most microalgal-bacterial functional interactions remain unknown, it is difficult to identify global trends in key functional types of bacteria associated with microalgal-bacterial communities. This urges the need for more functional studies to improve our understanding of microalgal-bacterial communities across the globe.

### Emerging microalgal-bacterial associations that can guide functional studies

An advantage of using a network approach is the ability to analyse large scale datasets to unravel previously unknown associations. We believe that some of the inferred co-occurrences in this study may help guide focused research to shed light on the functional nature of interactions. Therefore, we further explored these co-occurrences which were prominent due to their frequent observations and importance in modules based on topological properties.

For instance, uncultured *Deltaproteobacteria* order *NB1-j* was identified as consistently co-occurring with *Bacillariophyta* in marine environments. In one environment where the *NB1-j-Bacillariophyta* link was observed (Surface water samples from Western English Channel, EBI accession: ERP016541), *Bacillariophyta* was the only microalgal node directly interacting with *NB1-j*. In another environment (Seawater metagenome samples from Catlin Arctic survey (2010 expedition), EBI accession: ERP020022), *NB1-j* was identified as the hub node with the highest node degree (43), in a module consisting of 77 nodes and 817 edges. This *NB1-j* node also had the highest closeness centrality (0.69). Closeness centrality measures how close a node is to another node and helps to identify centrally positioned taxa in the network^24^. Other than this *NB1-j* node, there were 3 more *NB1-j* nodes in this community. All together these 4 *NB1-j* nodes were directly interacting (edges) with 57 nodes (out of 73 other nodes) of which 25 were representing microalgal nodes (Figure 3). Out of the 25 microalgal nodes, 13 represented the taxonomic class *Bacillariophyta*. Some of these *Bacillariophyta* were taxonomically identified at genus level as *Chaetoceros* and *Fragilariopsis*. Interestingly, except for one, all these Bacillariophyta nodes represented top hub nodes of the community with node degrees > 26. Most connected taxa are believed to have ecological relevance to the community as their removal causes the highest impact on many associations^74^.

**Figure 3:**
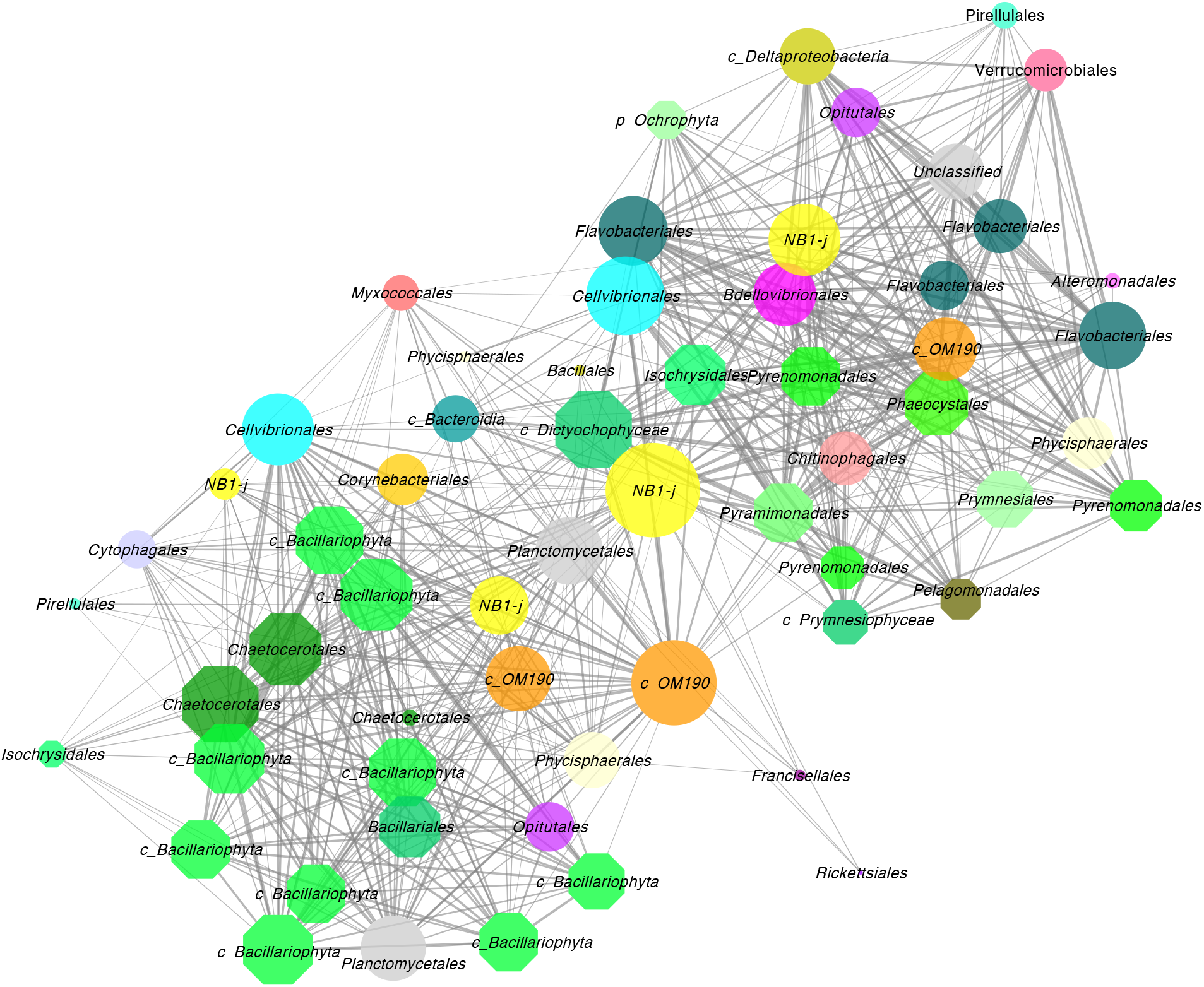
Interactions of the Deltaproteobacterial order *NB1-j* with microalgae in a marine environment. Figure shows the first neighbours (directly interacting taxa) including microalgal taxa co-occurring with Deltaproteobacterial order *NB1-j*. Nodes are labelled at the order level (If unclassified at the order level, higher taxonomic affiliations are provided). The edge width and node size are continuously mapped to edge weight (correlation strength) and node degree, respectively. Network image was generated using Cytoscape v3.8 (https://cytoscape.org/)

Functional roles of *NB1-j* in marine ecosystems are largely unknown. It is believed to be involved in hydrocarbon degradation^75^ and has been reported mostly from marine sediments ^70,76^, mud volcanoes^75^, sponges^77^ and cyanobacterial mats^78^. To the best of our knowledge, this *Deltaproteobacteria* group is not commonly reported in algal microbiomes. However, a recent study found *NB1-j* ASVs associated with the coral skeleton algal symbiont *Ostreobium*^79^. A predictive metagenomic approach based on sponge samples originating from reef sites in West Java (Indonesia) suggest that *NB1-j* may be involved in nitrogen metabolism^77^. It was found that *NB1-j* was responsible for the elevated predictive gene count corresponding to N-cycling genes such as those encoding nitric oxide reductase (*norB)*, nitrogenase (*nifD*) and hydroxylamine reductase. The available information indicates that the nitrogen cycling capacity of *NB1-j* may underly its association with algae, potentially facilitating the nitrogen needs of the algae while benefiting from algal organic carbon. However, a repertoire of interactions, as often expected from a keystone species, can be hypothesised between *NB1-j* and microalgae in a community.

*Bacillariophyta* (*Fragilariopsis, Chaetoceros* and many other diatoms unidentified at lower taxonomic levels) also showed frequent co-occurrences with an uncultured clade of *Planctomycetes, OM190* (SILVA taxonomy) in both freshwater and marine environments. In the Catlin Arctic survey samples (ERP02022), *OM190* demonstrated frequent associations with *Bacillariophyta*. Three *OM190* nodes were observed with high node degrees (29, 30 and 39) and these were directly connected to 59 nodes out of a total of 74 other nodes in the community. From the total of 35 microalgal nodes in this community, *OM190* were directly connected to 27 nodes out of which 15 nodes were represented by *Bacillariophyta* (Figure S3A). Similarly in a freshwater environment (Lake Superior, Michigan, EBI accession: ERP016492), *OM190* was identified as the 6^th^ most connected node with a node degree of 89 and was directly connected to 32 microalgal nodes including *Bacillariophyta* (Figure S3B). As mentioned previously, highly connected nodes act as hub nodes in the community. Multiple *OM190* acting as top hub nodes within their habitats indicate that they have high co-occurrences with both microalgae and bacteria.

*OM190* shows deep branching within the *Planctomycetes* group and is usually found in different environments such as soil and seawater. Members of this clade are usually considered to be associated with macroalgae^80^, such as red^81^ and brown algae^82^. Since *OM190* is yet uncultured, information on its metabolism is scarce. A metagenome-assembled-genome (MAG) for *OM190* (likely *OM190*) with a rich diversity of secondary metabolite potential has been reported^83^. The production of secondary metabolites by *OM190*, including antimicrobial compounds may be one of the underlying reasons for their association with algae as they could protect the alga from undesired microbes. In a recent study^84^, it was shown that diatoms produce fucose-containing sulfated polysaccharides (FCSP) which can be hypothesised as an energy source for *OM190*. Many *Planctomycetes* have abundant sulfatases, but these were not confirmed in the existing *OM190* MAG. In addition to their association with algae, *OM190* and *NB1-j* seem to have a close relationship with one another. They were recently reported as abundant co-occurring taxa in Beaufort Sea surface sediments^85^. The reasons for their direct associations are not known and, along with their respective relationships with algae, this direct interaction is an interesting topic for future investigation.

More generally speaking, *Planctomycetes* often associate with macroalgae and favour a biofilm lifestyle^82^. Protein clusters that may be involved in *Planctomycetes* symbiosis or biofilm maintenance have been reported^53^, and production of sulfated polysaccharides by the alga that serve as the substrate for the abundant sulfatases produced by the *Planctomycetes* contributes to the reasons for successful associations between them^53,80^. Although successful associations are known between macroalgae and *Planctomycetes*, interactions with microalgae are largely unknown. Apart from their associations with diatoms, our network analysis identified taxa representing *Planctomycetes* (c_*Phycisphaerae*, c_*Planctomycetacia*) co-occurring with a range of microalgal groups. In a previous study, *Planctomycetes* closely related to *Pirellula* were identified as one of the dominant lineages associated with diatom blooms^86^. Results of our network analysis demonstrate that associations between *Planctomycetes* and microalgae may be as common as those they maintain with macroalgae.

*Verrucomicrobia* were also consistently associated with an array of microalgae in fresh and marine environments indicating that they may be common associates of microalgae. Here in our study, all the Verrucomicrobial members co-occurring with microalgae were represented by the taxonomic class *Verrucomicrobiae*. Among the genus-specific associations revealed in our network analysis is that of the Verrucomicrobial genus *Lentimonas* and microalgal genera *Pyramimonas* (c*_Prasinophyceae*), *Phaeocystis, Tisochrysis, Haptolina*, and *Chrysochromulina* (c_*Prymnesiophyceae*). A recent study showed hundreds of *Lentimonas* enzymes able to digest brown algal fucoidan^87^. Although fucose-containing sulphated polysaccharides were regarded as a macroalgal polysaccharide, microalgae such as diatoms have the potential to produce them^84^. We speculate that other microalgae might also produce this sulfated polysaccharide, or that members of *Lentimonas* may have a broader palate than just FCSP. Although widely distributed, the functional roles of *Verrucomicrobia* in aquatic ecosystems are not well understood due to the lack of cultured strains^88^. Some members of *Verrucomicrobia* have been shown to consume algal extracellular polymeric substances^69,89,90^. However, other factors contributing to their associations are not well understood.

## Conclusion

In summary, our study illustrates the promise of using 16S rRNA gene-based co-occurrence networks as a hypothesis-generating framework to guide focused research and speculate on the functional nature of potential interactions. By studying multiple environments based on public datasets, we provided an overview of microalgal-bacterial communities in aquatic ecosystems from a network perspective. This identified a range of associations including previously unknown links that can set the stage for more focused research in the future.

## Declarations

### Ethics approval and consent to participate

Not applicable

### Consent for publication

Not applicable

### Availability of data and material

No data was generated in this study and all the data analysed are publicly available. The EMP datasets analysed are available in European Nucleotide Archive and EMP Qiita portal under the Accession numbers and Qiita ids, respectively, provided in Supplementary material TableS1. FastSpar script used for the correlation analysis, R script used in generating networks and the networks files generated in Cytoscape can be found at https://melbourne.figshare.com/projects/Unravelling_microalgal-bacterial_interactions_in_aquatic_ecosystems_through_16S_rRNA_gene-based_co-occurrence_networks/140071

### Authors’ contribution

UP, HV and AW formulated the project. UP analysed the data and wrote the manuscript with inputs from all the authors. HV, AW and KT contributed ideas and helped with analysis. All authors read and approved the final version of the manuscript.

## Acknowledgements

Not applicable

## Supplementary Data

**Figure S1: Map indicating the sample collection sites for each EMP project**. Sample collection locations for each EMP project are shown on the world map. Geographic coordinates were drawn in R studio (v1.4.1106) using packages rworldmap v1.3-6 (https://rdrr.io/cran/rworldmap/).

**Figure S2: Major bacterial Phyla associated with microalgae in freshwater and marine environments**

Bars represent the number of bacterial nodes affiliated with each phylum. The total number of bacterial nodes identified to be co-occurring with microalgae in marine environments were 76 and freshwater environments were 311.

**Figure S3: *Planctomycetes* class *OM190* interacting with microalgal taxa in a marine (A) and freshwater (B) environment**.

Networks (A, B) created from a microalgal-bacterial module by selecting the first neighbours of the *OM190* to showcase the complex interactions with the neighbouring microalgal taxa. The edge width and node size are continuously mapped to edge weight and node degree. Marine and Freshwater environments represent EMP project ERP020022 and ERP016492, respectively. Nodes labelled at order level (If unclassified at the order level, higher taxonomic affiliations are provided). Network image was generated using Cytoscape v3.8 (https://cytoscape.org/).

**Table S1: Information on selected EMP studies**. Details on Qiita study title, study ids and ENA project accession number.

**Table S2: Correlations recovered in marine environments between microalgae and bacteria**. Summary of significant co-occurrences recovered from the marine environments. Full taxonomic affiliations of the microalgal and bacterial nodes are provided.

**Table S3: Correlations recovered in freshwater environments between microalgae and bacteria**. Summary of significant co-occurrences recovered from the freshwater environments. Full taxonomic affiliations of the microalgal and bacterial nodes are provided.

