## Supplementary figures and images for "Unravelling microalgal-bacterial interactions in aquatic ecosystems through 16S rRNA gene-based co-occurrence networks"

### Figure S1

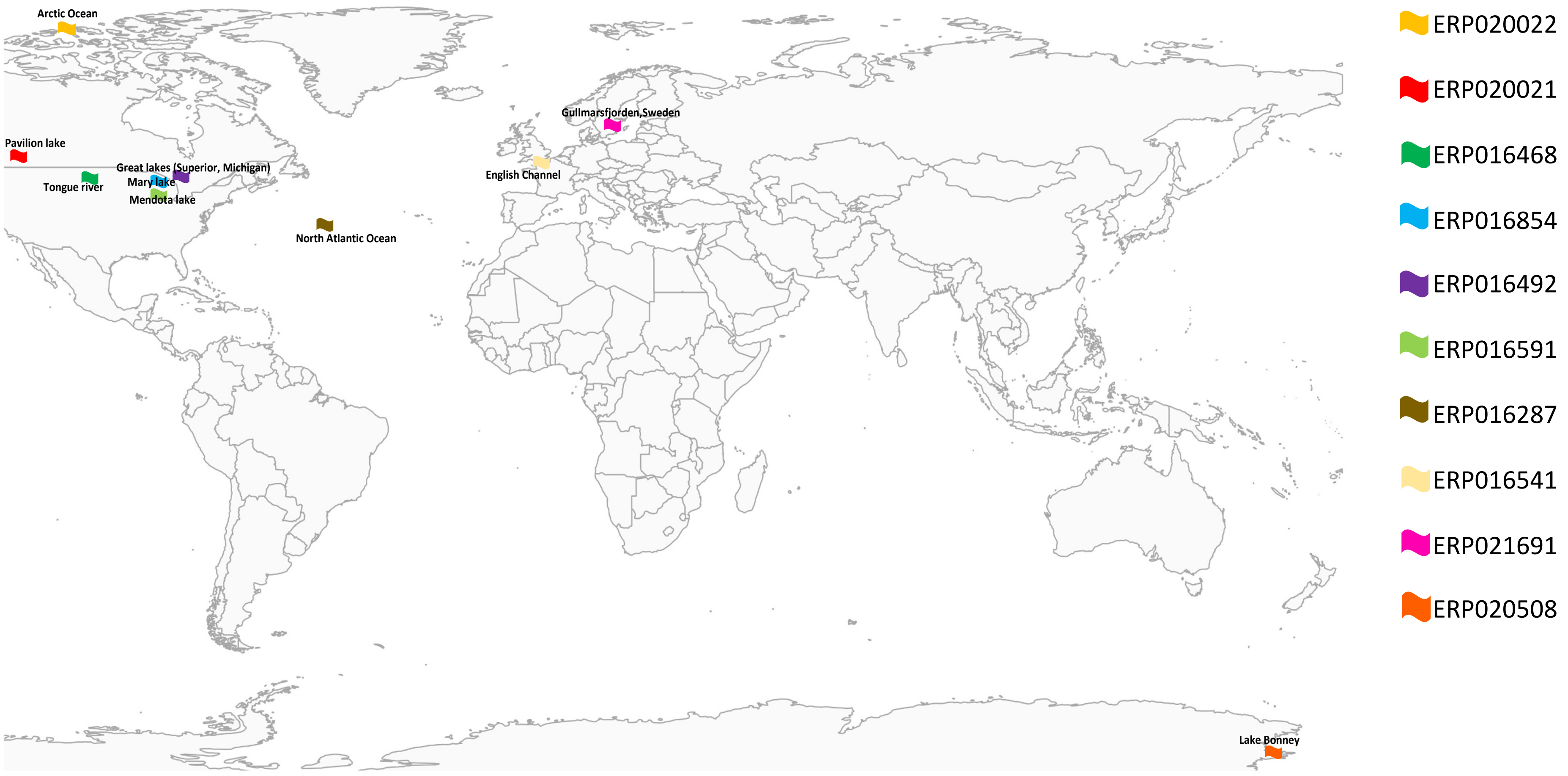

### Figure S2

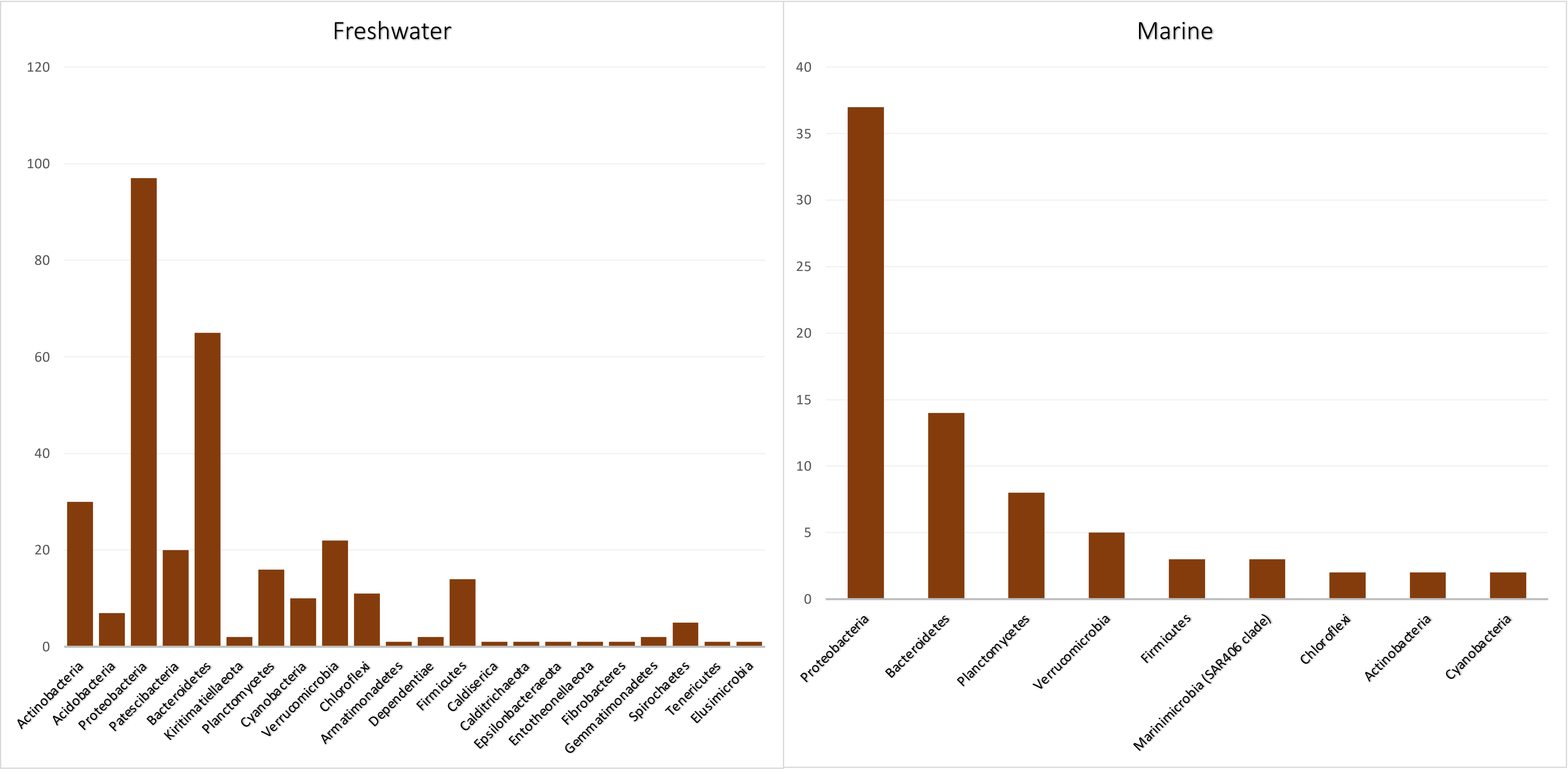

### Figure S3

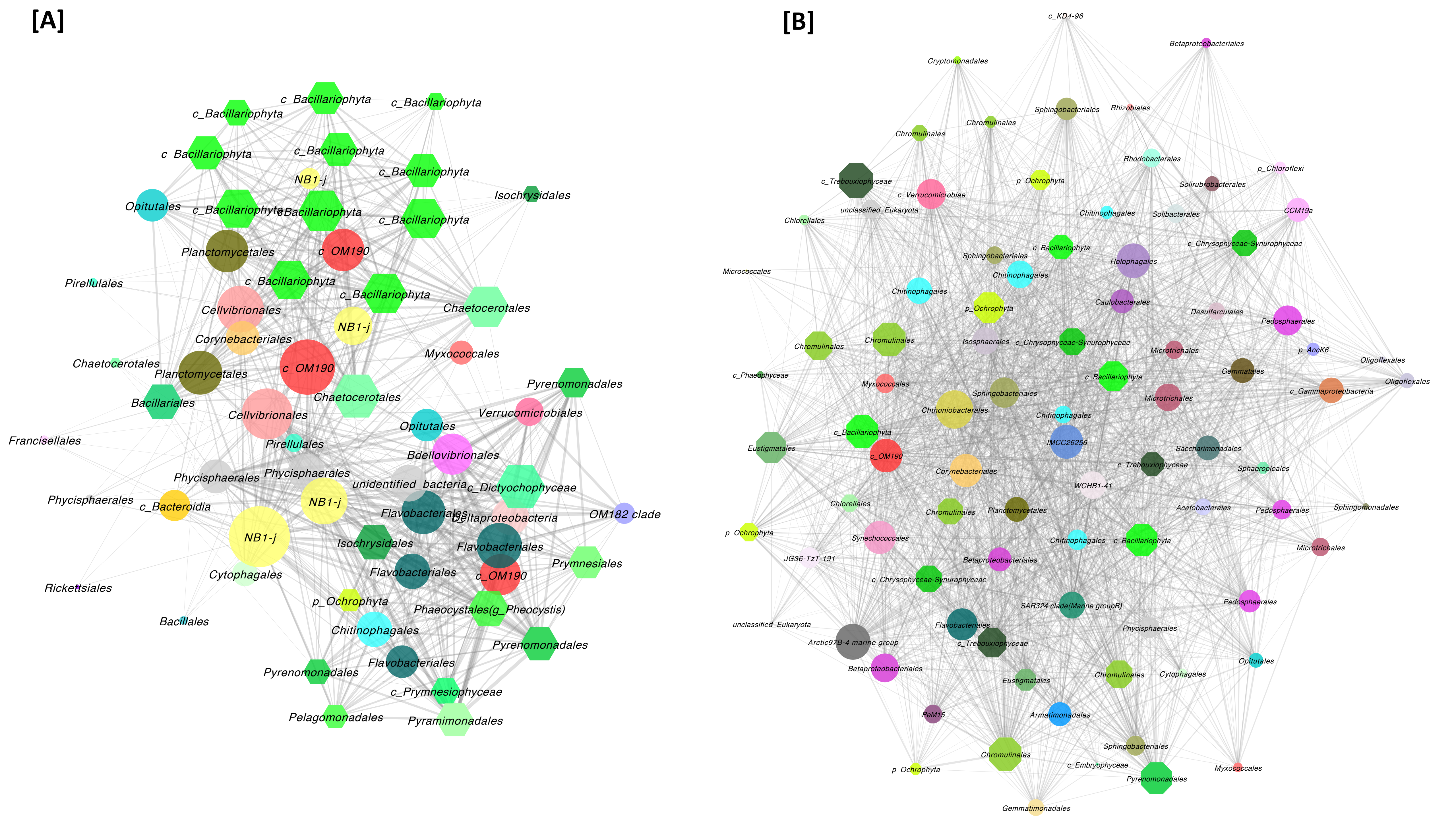
